# The Axenfeld-Rieger syndrome gene *FOXC1* contributes to left-right patterning

**DOI:** 10.1101/2020.05.28.120915

**Authors:** Paul W. Chrystal, Curtis R. French, Francesca Jean, Serhiy Havrylov, Suey van Baarle, Ann-Marie Peturson, Pengfei Xu, J. Gage Crump, David B. Pilgrim, Ordan J. Lehmann, Andrew J. Waskiewicz

**Affiliations:** Department of Medical Genetics, University of Alberta, Edmonton, Alberta, T6G 2H7, Canada; Department of Ophthalmology, University of Alberta, Edmonton, Alberta, T6G 2H7, Canada; Department of Biological Sciences, University of Alberta, Edmonton, Alberta, T6G 2E9, Canada; Faculty of Medicine, Memorial University of Newfoundland, St John’s, Newfoundland, A1B 3V6, Canada; Department of Stem Cell Biology and Regenerative Medicine, University of Southern California, Keck School of Medicine, Los Angeles, CA 90033, USA; Women & Children’s Health Research Institute, University of Alberta, Edmonton, T6G 1C9, Canada

**Author notes:** Co-corresponding authors, Correspondence to: Andrew J. Waskiewicz, Department of Biological Sciences, CW405 Biological Sciences Building, 11455 Saskatchewan Dr., University of Alberta, Edmonton, AB T6G 2E9, Canada., Ordan J. Lehmann, Departments of Ophthalmology and Medical Genetics, 829 Medical Sciences Building, University of Alberta, Edmonton, AB T6G 2H7, Canada.

**Keywords:** FOXC1, Axenfeld-Rieger syndrome, left-right patterning, zebrafish, Lefty

## Abstract

Normal body situs requires precise spatiotemporal expression of the *Nodal-Lefty-Pitx2* cascade in the lateral plate mesoderm. The ultimate output of this patterning is establishment of the left-right axis, which provides vital cues for correct organ formation and function. Mutations, deletions and duplications in *PITX2* and *FOXC1* lead to the rare genetic disease Axenfeld-Rieger syndrome (ARS). While situs defects are not a recognised feature of ARS, partial penetrance of cardiac septal defects and valve incompetence is observed; both of these congenital heart defects (CHDs) also occur following disruption of left-right patterning. Here we investigated whether *foxc1* genes have a critical role in specifying organ situs. We demonstrate that CRISPR/Cas9 generated mutants for the zebrafish paralogs *foxc1a* and *foxc1b* recapitulate ARS phenotypes including craniofacial dysmorphism, hydrocephalus and intracranial haemorrhage. Furthermore, *foxc1a*^-/-^; *foxc1b*^-/-^ mutant animals display cardiac and gut situs defects. Modelling *FOXC1* duplication by transient mRNA overexpression revealed that increased *foxc1* dosage also results in organ situs defects. Analysis of known left-right patterning genes revealed a loss in expression of the *NODAL* antagonist *lefty2* in the lateral plate mesoderm. Consistently, *LEFTY2* mutations are known to cause human cardiac situs defects. Our data reveal a novel role for the forkhead-box transcription factor *foxc1* in patterning of the left-right axis, and provide a plausible mechanism for the incidence of congenital heart defects in Axenfeld-Rieger syndrome patients.

**Author Summary:** This manuscript investigates the functional consequences of abrogating the activity of Foxc1 (Forkhead Box C1). We demonstrate that loss of zebrafish *foxc1a* and *foxc1b* results in phenotypes that resemble human patients with deletions in the *FOXC1* locus. Notably, such phenotypes include alterations to the morphology of the heart. Investigations into the mechanisms underlying this phenotype led to the discovery that Foxc1 functions as a regulator of left-right patterning. Most components of left-right specification function normally in *foxc1a/b* mutants, but there is a pronounced loss of *lefty2*, a known inhibitor of Nodal signaling. This supports a model in which Foxc1 regulates situs of the heart via the regulation of Lefty2.

## Introduction

Establishment of left-right asymmetry represents a fundamental step in embryonic development. Despite substantial progress elucidating a proportion of the core players, the mechanisms remain incompletely defined. Consequently, syndromes where organs are aberrantly positioned are of particular interest to geneticists and developmental biologists. In humans, the heart’s normal anatomical position is left of midline, with a larger left ventricle designed for systemic circulation. The right lung has three lobes, while the left lung has two lobes and contains an indentation, the cardiac notch, allowing space for the heart. Furthermore, the stomach and liver are positioned left and right of the midline, respectively. This normal arrangement is called situs solitus, while the complete reversal of normal organ situs (termed situs inversus) is surprisingly well tolerated (1). Far more deleterious are partial situs defects, collectively known as heterotaxy (2), characterized by mis-patterning of visceral organs along the left-right axis. These are associated with congenital diseases of the heart, lungs, spleen, stomach and liver (3–5) that may be particularly challenging to treat. Intriguingly, many heterotaxy-associated genes also cause isolated congenital heart defects (CHDs) (6), suggesting that a proportion of idiopathic CHDs may reflect unrecognized situs defects (2, 7–9).

In vertebrates, the breaking of left-right symmetry is established around a structure known as the left-right organizer (LRO). In mouse, zebrafish, frog and humans, asymmetric fluid flow, generated by motile monocilia projecting into the extracellular fluid of the LRO, leads to asymmetric gene expression patterns around the LRO (10). Consequently, loss of flow or the motile cilia results in situs defects in animal models and humans (11–13). In many vertebrates the output of the LRO first manifests as decreased expression of *Dand5* (also known as *charon* in zebrafish or *Cerl2* in mouse), which normally represses the TGF-β family member *Nodal*, a secreted morphogen that is transiently expressed on the left side of the embryo (14, 15). *Nodal* upregulates its own transcription, as well as transcription of the homeobox domain transcription factor *Pitx2* (16). Despite asymmetric *Nodal* expression lasting only a matter of hours, murine *Pitx2* expression persists in the left lung and cardiac tissue throughout organogenesis and into adulthood (16–21). Pitx2 plays a critical role in the establishment of left-right asymmetry and homozygous murine mutants display pronounced phenotypes. For example, at E12.5, murine *Pitx2* homozygotes display cardiac situs defects and right pulmonary isomerism (identical lobar anatomy of the left and right lungs) (19), while second heart field-specific *Pitx2* mutations cause severe cardiac outflow tract defects (22). In patients, heterozygous *PITX2* mutations cause Axenfeld-Rieger syndrome (ARS), and despite multi-organ involvement, organ situs defects have not been observed (23–25). This likely reflects patients’ heterozygous variants, compared with homozygous deletion of *Pitx2*, that can be achieved either globally or in the secondary heart field of murine models (19, 22).

Precise control of left-right patterning relies on the establishment of a signaling barrier to separate left– and right–specific gene expression programs. Critical components of the midline barrier are the TGF-β family members Lefty1/2 (26, 27) that are expressed in the midline and on the left side diffuse to the midline and right side to inhibit Nodal signaling. Although both genes are expressed in the same region, the increased diffusion of Lefty proteins (relative to Nodal) limits Nodal target gene activation on the right side of the embryo (28). Consequently, mice with a deletion in the *Lefty2* enhancer that is activated by Nodal, display left isomerism (29) while variants in *LEFTY1/2* are associated with congenital heart defects (30). Illustrating significant additional complexities in the control of left-right patterning, pathways initiated on the right side of the embryo have also been shown to contribute to organ laterality, as demonstrated by BMP-dependent activation of Prrx1a in cardiac laterality and the role of hyaluronan in determining midgut laterality (31, 32).

Axenfeld-Rieger syndrome (ARS) is an autosomal dominant condition caused by mutation and copy number variation of *PITX2* and *FOXC1* (24, 33–35). The ARS phenotypic spectrum includes ocular anterior segment dysgenesis, early-onset glaucoma, craniofacial dysmorphism, cerebral small vessel disease, cerebellar vermis hypoplasia, and hydrocephalus (36–41). Despite no reported association with laterality defects, congenital heart defects are present in ARS and these include atrial and ventricular septal defects, valve stenosis and persistent truncus arteriosus (24, 42–46) particularly associated with *FOXC1* mutations (23, 24, 47, 48). Murine *Foxc1* mutants also exhibit CHDs, but no analysis of L-R patterning has been completed (49).

To test the hypothesis that Foxc1 is a regulator of L-R patterning, we mutated the two zebrafish paralogs via CRISPR/Cas9 editing. In addition to replicating many of the ARS-associated phenotypes observed with human *FOXC1* mutation, this strategy yielded evidence for a requirement for zebrafish *foxc1a/b* in establishing cardiac and gut laterality. Analysis of these zebrafish mutants also established that the expression of *lefty2* was altered. Our data thus provide the first evidence for a contribution by Foxc1 to left-right patterning.

## Results

### Generation of foxc1a and foxc1b zebrafish mutants

Zebrafish possess two orthologs of human *FOXC1* (*foxc1a* and *foxc1b*) but lack a *FOXC2* ortholog (50). While it has been suggested that zebrafish *foxc1b* may represent a functional ortholog of human *FOXC2* (51), the amino acid sequences of Foxc1a and Foxc1b are more similar to human FOXC1 (identities: 74% and 68%, respectively) than FOXC2 (55% and 53%) (Table S1), with equivalent results when comparing the DNA-binding, *forkhead* domains (FOXC1: 97% and 97%, FOXC2: 93% and 93%). Since the zebrafish paralogs possess conserved synteny with human *FOXC1* (Fig. S1), as well as similar protein sequences and overlapping expression (50, 51), we anticipated compensatory activity with loss of a single paralog. To address this, we generated mutations in both *foxc1a* and *foxc1b* using CRISPR/Cas9 mutagenesis. The *foxc1a*^UA1017^ allele is a 7 nucleotide deletion that is predicted to cause a premature stop codon after 39 amino acids (c.29_35del; p.Pro10Serfs*39). The *foxc1b*^UA1018^ allele is a 40 nucleotide deletion that results in a premature stop codon after 28 amino acids (c.57_96del; p.Ile19Metfs*28) (Fig. 1A-C). Both changes (hereafter referred to as *foxc1a*^-/-^ and *foxc1b*^-/-^) lie upstream of the forkhead domain, the alleles are putative nulls due to predicted loss of >90% of the encoded protein (Fig. 1B). Analysis of *foxc1a/b* gene expression within the mutants revealed no evidence for nonsense mediated decay, which is consistent with other findings for single exon loci (Fig. S2).

**Fig. 1.**
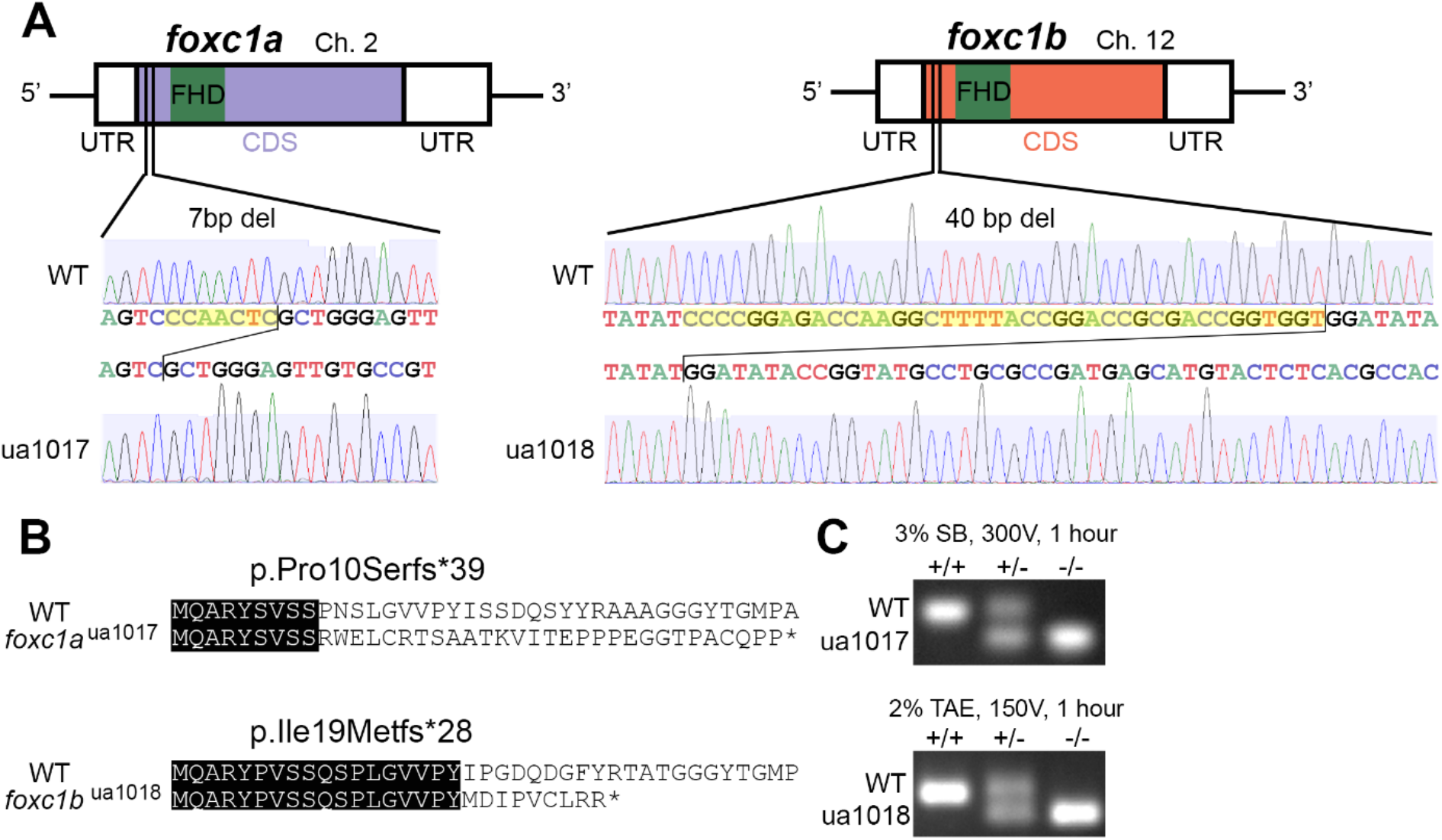
Allelic consequence of *foxc1*-targeted CRISPR/Cas9 mutagenesis. (**A**) Schematic representation of *foxc1a*^ua1017^ and *foxc1b*^ua1018^ mutations, with 7 and 40 base pair deletions (yellow highlighting) upstream of the DNA-binding, *Forkhead* domain. (**B**) The predicted sequences of the first 40 amino acids translated from wildtype (WT) and mutant proteins with the sequence identity (black highlighting). The *foxc1a*^ua1017^ allele produces a truncated 39-residue protein with loss of sequence homology from amino acid 10. The *foxc1b*^ua1018^ allele produces a truncated 28-residue protein with loss of sequence homology from amino acid 19. (**C**) PCR genotyping from gDNA template resolves the respective deletions in *foxc1a* and *foxc1b*. (FHD, *Forkhead* domain; UTR, untranslated region; CDS, coding sequence).

### foxc1a single and double mutants display gross developmental defects

*foxc1a*^+/-^ zebrafish are viable and fertile as heterozygotes and generate Mendelian ratios of offspring when incrossed. However, *foxc1a*^-/-^ homozygotes do not live beyond 7 days post fertilization (dpf) and display obvious developmental defects by 96 hours post fertilisation (hpf; Fig 2A-F). Blood flow is observed until 72 hpf (data not shown) when there is pericardial oedema in 86% of embryos (Fig. 2C), that becomes more severe over time. Zebrafish *foxc1a*^-/-^ homozygotes also display microphthalmia (Fig. 2B, C) and intracranial haemorrhage (Fig. 2D) at 72 hpf. *foxc1a*^-/-^ homozygotes display craniofacial dysmorphism (59%; Fig. S3), which has been reported previously (52).

**Fig. 2.**
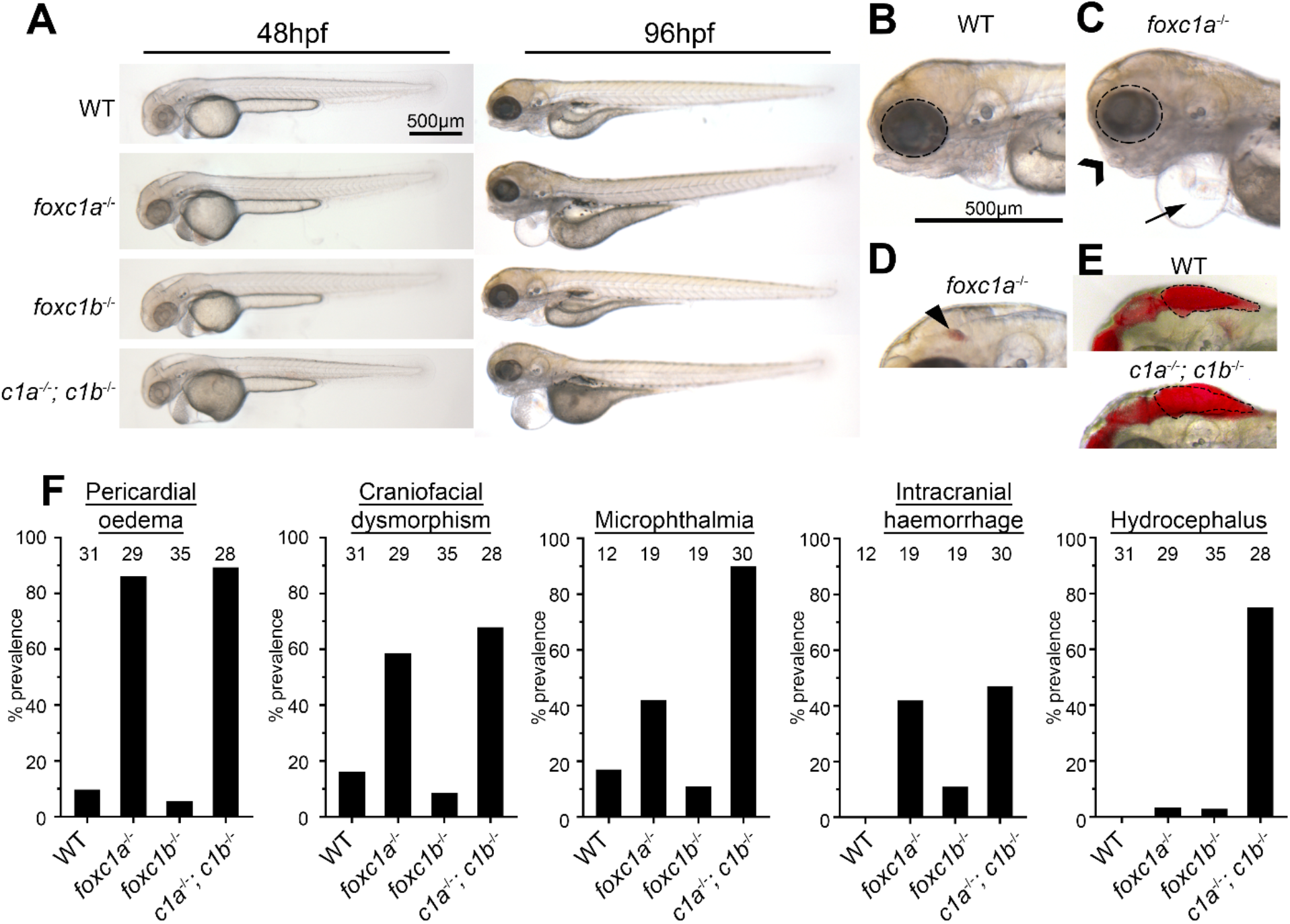
*foxc1* mutants exhibit multiple developmental defects. (**A**) *foxc1* single mutants are largely indistinguishable from WT (wild type) controls at 48 hpf (left panels) whereas *foxc1a*^-/-^; *foxc1b*^-/-^ double homozygotes display hydrocephalus and oedema. By 96 hpf no changes are observed in *foxc1b*^-/-^ homozygotes, however, *foxc1a*^-/-^ homozygotes and *foxc1a*^-/-^; *foxc1b*^-/-^ double homozygotes display pericardial oedema (compare **B** and **C**, as highlighted by arrow), microphthalmia (dotted circle), and craniofacial dysmorphism (chevron). At this stage a subset of mutants also displayed intracranial haemorrhage (arrowhead in panel **D**) with *foxc1a*^-/-^ homozygotes having greater frequency than *foxc1b*^-/-^ homozygotes (42% vs 11%, respectively), and (**E**) *foxc1a*^-/-^; *foxc1b*^-/-^ double homozygotes present with hydrocephalus. (**F**) Quantification reveals that these defects are incompletely penetrant and generally more prevalent in double than single homozygotes (number of embryos analyzed is shown above each bar). In the case of hydrocephalus, only the double homozygotes display an appreciable frequency of this phenotype.

Zebrafish *foxc1b*^-/-^ homozygotes are viable as larvae, survive to adulthood and are fertile with no observable phenotype (results consistent with other studies (51, 52)). Incrossing two *foxc1b*^-/-^ homozygous adults to generate maternal zygotic *foxc1b*^-/-^ homozygote offspring also generated viable embryos indistinguishable from wildtype controls. Prior research on the zebrafish *foxc1b*^-/-^ determined that it is a null allele (53). To investigate the potential for genetic compensation, a double *foxc1a/foxc1b* mutant line was generated. Both double heterozygotes (*foxc1a*^+/-^; *foxc1b*^+/-^) and fish with 3 mutant alleles (*foxc1a*^+/-^; *foxc1b*^-/-^) were viable as adults, fertile, and exhibited no phenotypes. Crossing two double heterozygotes generated double mutant homozygotes (*foxc1a*^-/-^; *foxc1b*^-/-^) at expected Mendelian ratios, although developmental deformities were severe, especially with regards to microphthalmia and hydrocephalus (Fig. 2E-F). 89% of *foxc1a*^-/-^; *foxc1b*^-/-^ double homozygotes displayed pericardial oedema at 72 hpf, at which time blood flow had ceased. Craniofacial dysmorphism was observed in 68% of embryos and was more severe than single *foxc1a*^-/-^ homozygotes. Additionally, 75% of *foxc1a*^-/-^; *foxc1b*^-/-^ double homozygotes displayed hydrocephalus, which was absent in single mutants (Fig. 2E-F). These data demonstrate that loss of zebrafish *foxc1a/b* paralogs is generally more severe than single homozygosity, and supports the hypothesis of genetic buffering by the two paralogs.

### Alterations to foxc1 gene dosage cause visceral organ situs defects

Due to the progressive pericardial oedema and loss of blood flow observed from 72 hpf in *foxc1a* homozygotes, we next examined cardiac development in more detail. Loss of *foxc1* resulted in significantly fewer embryos with normal D-looped hearts, and increased the prevalence of abnormal O-looped (unlooped) and L-looped (situs inversus) hearts (Fig. 3A). Homozygous *foxc1a* mutant hearts failed to loop in 31% of those examined (n=32, P=0.016). Homozygous *foxc1b* mutants also displayed a significant divergence from normal cardiac looping (n=39, P=0.036) with 23% O-loop and 5% L-looped hearts. Double *foxc1a*^-/-^; *foxc1b*^-/-^ loss resulted in only 38% of embryos developing D-loops with the remaining 62% producing O-loops (n=29, P<0.001), consistent with overlapping functions of *foxc1* paralogs in cardiac situs phenotypes.

**Fig. 3.**
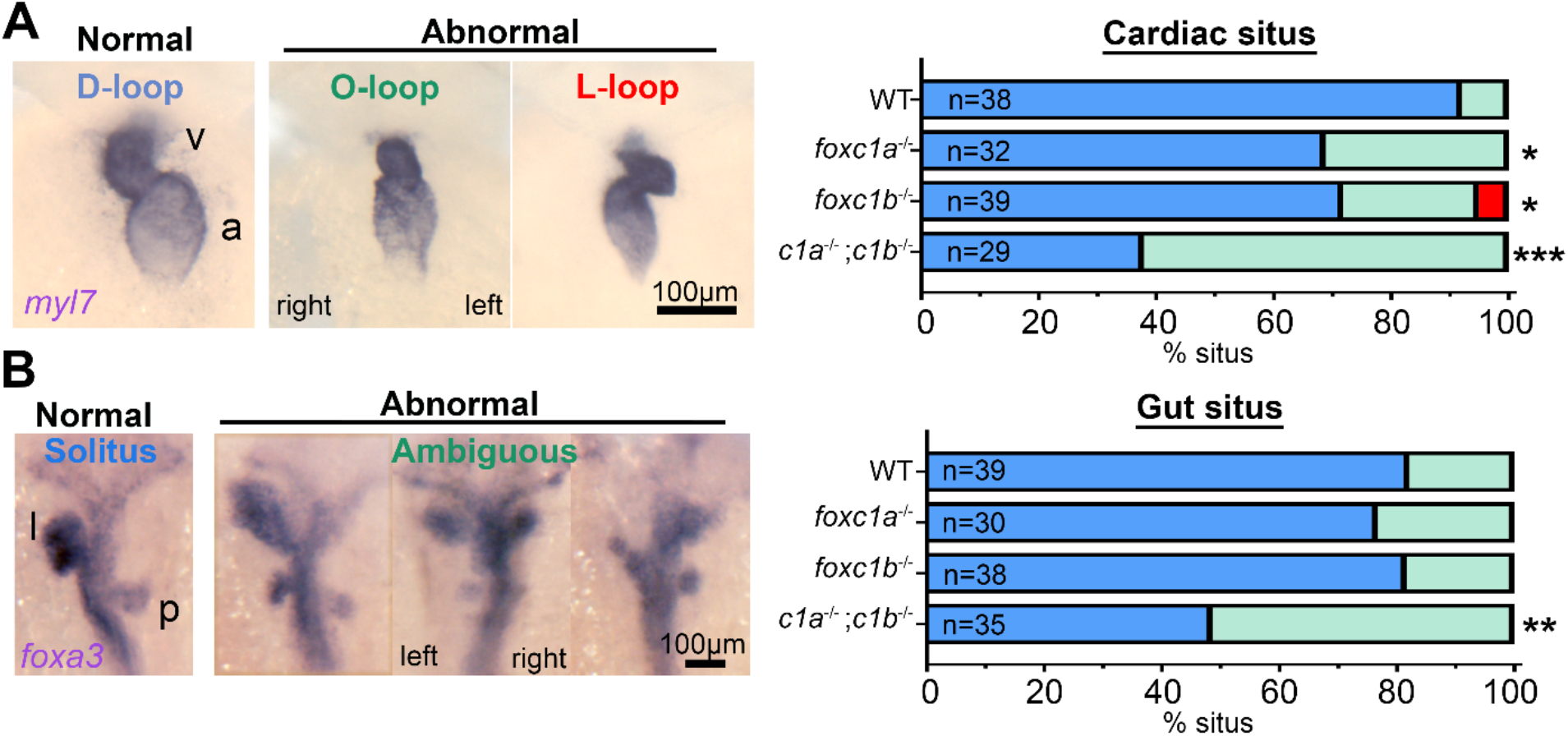
Decreased and increased dosage of *foxc1* result in multi-organ situs defects. (**A**) *In situ* hybridisation with *myl7* at 48 hpf revealed aberrant cardiac situs [O-loop (green); L-loop (red)] compared with normal D-loop (blue). The prevalence of aberrant situs is increased in *foxc1a, foxc1b*, and double homozygotes (*P=0.016, *0.036, ***<0.001, respectively Fisher’s exact test) when compared to WT siblings. (**B**) At the same stage, the normal arrangement (solitus, blue) of the gut is left-sided liver (l) and right-sided pancreas (p), however the incidence of abnormal (ambiguous, green) gut situs was significantly greater in double homozygotes (**P=0.003, Fisher’s exact test).

To investigate whether the situs defects were cardiac specific or systemic, gut situs was assessed using the liver and pancreas expression domain of *foxa3*. Although neither *foxc1a*^-/-^ or *foxc1b*^-/-^ homozygotes had altered gut situs compared to controls, 51% of *foxc1a*^-/-^; *foxc1b*^-/-^ double homozygote embryos developed abnormal gut situs. In those embryos, the most common observation was isomerism of the liver and or pancreas (Fig. 3B). These results suggest that cardiac situs is more sensitive to loss of *foxc1* than gut situs.

Because *FOXC1* function is exquisitely sensitive to gene dosage, with both gene duplication and deletion causing disease (23), we examined whether mRNA overexpression of *foxc1* induces cardiac situs defects (Fig. 4). Injection of 75 pg of either *foxc1a* or *foxc1b* mRNA resulted in divergence from normal D-looped hearts observed in 45% of *foxc1a* and 38% of *foxc1b* mRNA injected embryos, prevalences greatly increased from the mCherry mRNA control (Fig. 4A-C). Such cardiac situs defects were observed in a dose-dependent manner (Fig. 4D) and hydrocephalus commonly arose at 72 hpf in embryos overexpressing either *foxc1a* or *foxc1b* (Fig. S4). Gut situs was similarly affected (Fig. 4E), with comparable prevalence of abnormal gut situs observed (*foxc1a* overexpression: 53%, *foxc1b* 31%, mCherry control 3 %, P=0.02).

**Fig. 4.**
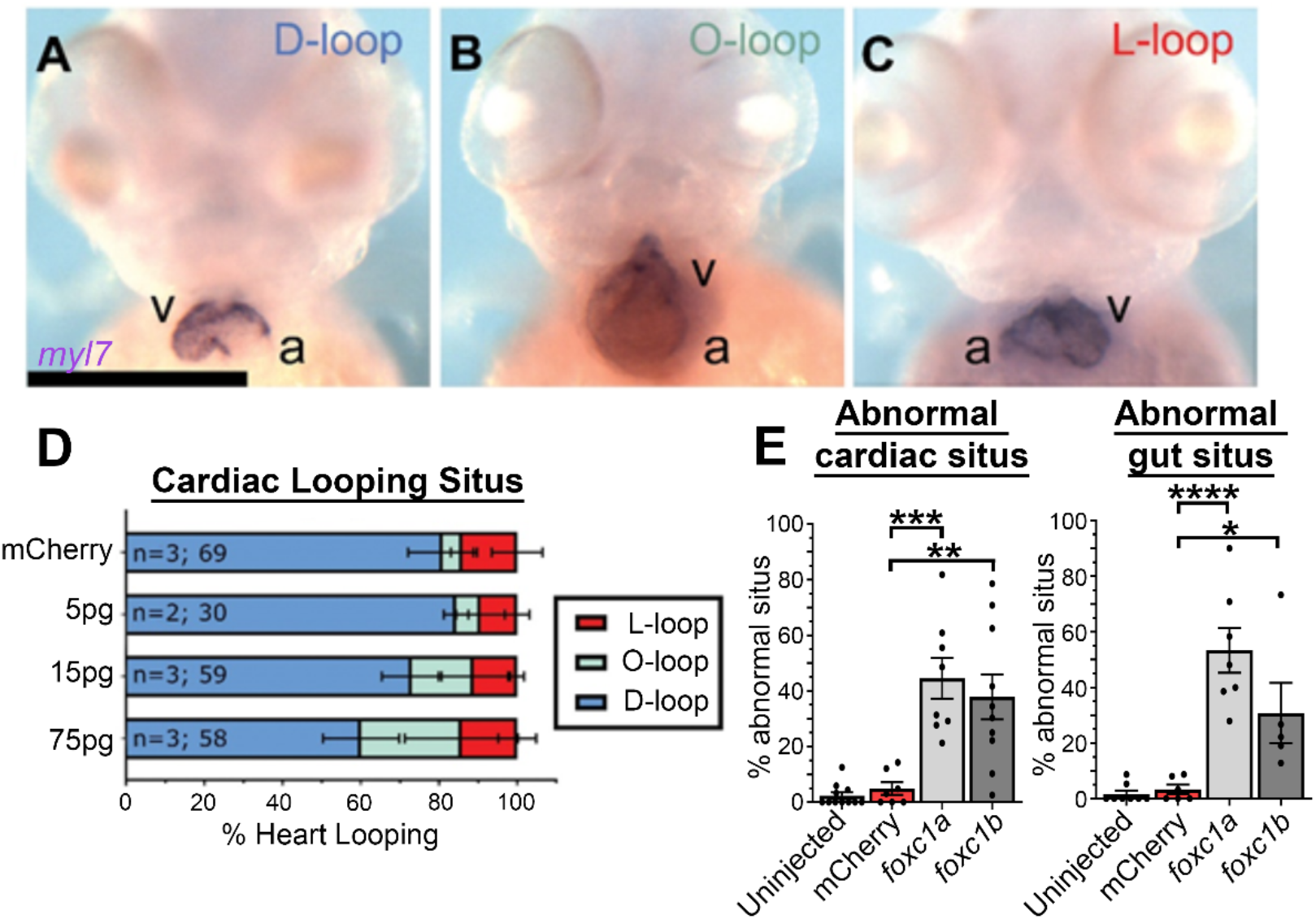
*foxc1a* mRNA overexpression causes cardiac situs defects in a dose-dependent manner. (**A-C**) *myl7* in situ hybridisation of 75 pg *foxc1a* mRNA injected embryos at 48 hpf, with representative images of the three cardiac looping morphologies provided (v, ventricle; a, atrium). (**D**) Quantification of cardiac situs in control and *foxc1a* mRNA injected embryos revealed an increasing prevalence of anomalous cardiac looping with increasing amounts of *foxc1a* mRNA. (E) Quantification of embryo situs defects in embryos injected with 75 pg *foxc1a* mRNA. Statistical significance was apparent in comparisons between *foxc1a/b* and *mCherry* controls (*P<0.05, **P<0.01, ***P<0.001, ****P<0.0001) If you want exact values then: Cardiac: c1a = 0.0002, c1b = 0.0012. Gut: c1a = <0.0001 (graphpad doesn’t calculate if <0.0001), clb = 0.0205. one-way ANOVA and Dunnette’s Posthoc test.

Although these data support a requirement for *foxc1* in normal organ situs determination in zebrafish and were supported by morpholino knockdown (Fig. S5), this finding ran counter to conventional understanding of Foxc1’s roles. Therefore, we validated these results by comparing phenotypes with those observed in independently generated zebrafish alleles (52). These *foxc1a*^-/-^; *foxc1b*^-/-^ double homozygotes exhibited a similar prevalence of cardiac and gut situs defects that were absent from wild type siblings (Fig. S6).

### Loss of foxc1a/b does not disrupt LRO fluid flow

Several vertebrate species, including zebrafish, possess a left-right organiser (LRO) where *foxj1*-dependent, motile cilia generate leftward fluid flow to create left-right symmetry. Accordingly, we next examined the LRO cilia in single and double *foxc1* mutants to determine if alterations were present. Alterations to axonemal length of *foxc1a*^-/-^ and double *foxc1a*^-/-^; *foxc1b*^-/-^ homozygotes (Fig. 5A; each 82% of WT) did not reach statistical significance (P=0.279). Fluorescent bead tracking experiments in *foxc1a*^+/-^ heterozygous incrosses revealed comparable ciliary-driven counter-clockwise flow in wild type embryos, *foxc1a*^-/-^ homozygotes, and *foxc1ab* morphants (Fig. 5B, B’, B”). Together, these data suggest that loss of *foxc1a/b* does not appreciably alter LRO ciliary function.

**Fig. 5.**
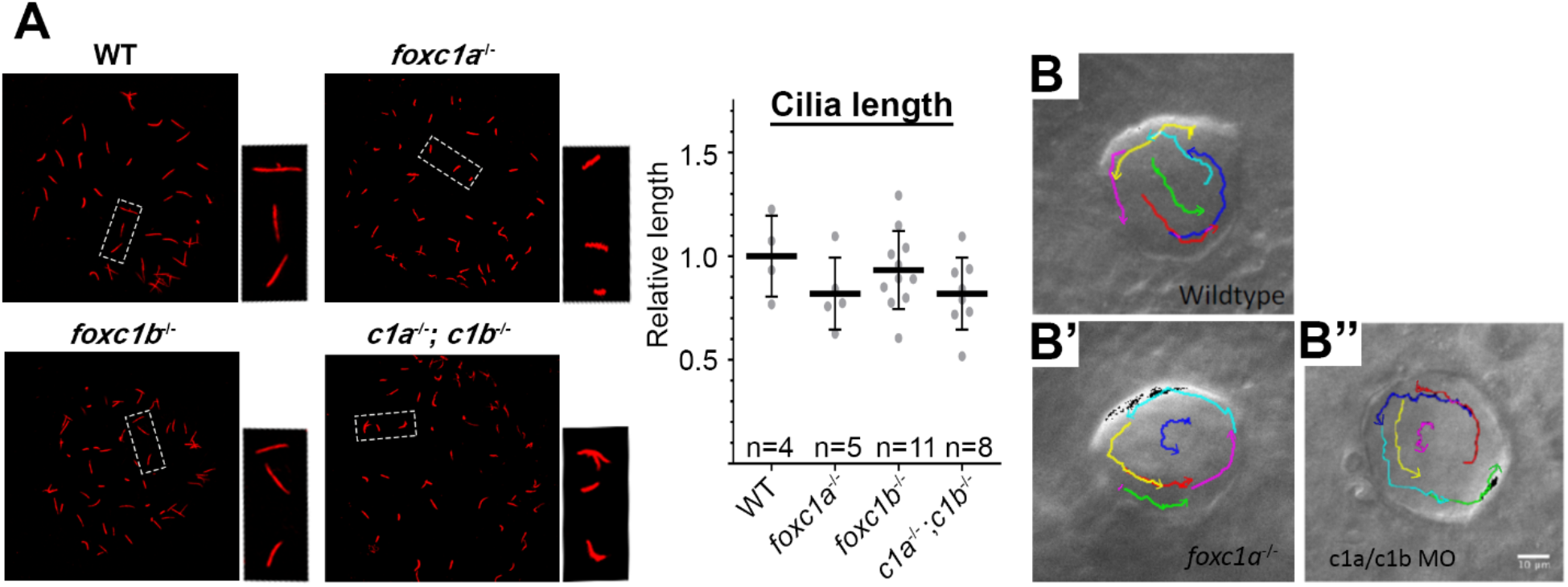
Loss of *foxc1* does not significantly change cilia length or Left-Right Organiser flow. (**A**) Acetylated alpha-tubulin immunostaining of LRO cilia revealed that average cilia length was not significantly altered in *foxc1* mutants (*foxc1a*^-/-^, 82%; *foxc1b*^-/-^, 93%; *foxc1a*^-/-^ *foxc1b*^-/-^, 82% of WT length; P= 0.286, ANOVA; 4-11 embryos imaged per condition). Tracking of fluorescent bead flow in the LRO revealed unchanged counter-clockwise flow between WT (**B**), *foxc1a*^-/-^ homozygotes (**B’**) and *foxc1a/foxc1b* morphants (**B”**).

### foxc1 mutants have loss of lefty2 expression in the lateral plate mesoderm

Since no changes at the level of the LRO were resolved, we next examined gene expression downstream of the LRO. The early L-R patterning gene *southpaw* (*spaw*), equivalent of mammalian *Nodal*, did not show altered expression patterns in *foxc1* mutants (Fig. 6A). In contrast, the Nodal antagonist *lefty2* was altered in the lateral plate mesoderm (Fig 6B): 33% of *foxc1a*^-/-^, 43% of *foxc1b*^-/-^, and 38% of *foxc1a*^-/-^; *foxc1b*^-/-^ double homozygotes exhibited normal left-sided expression (Fig. 6B). Similar results were observed with the independently generated *foxc1a*^-/-^; *foxc1b*^-/-^ double homozygotes (Fig. S6). Overexpression of both paralogs was next performed to determine if *lefty2* expression was affected by increased *foxc1* levels, revealing the loss of left sided *lefty2* expression in approximately 83% of *foxc1a* and 77% of *foxc1b* overexpressing embryos, prevalence significantly altered when compared with mCherry control (P=0.0007, 0.003 respectively) (Fig. 6C). Together, these data demonstrate that increased, and decreased *foxc1* expression, can perturb *lefty2* patterning.

**Fig. 6.**
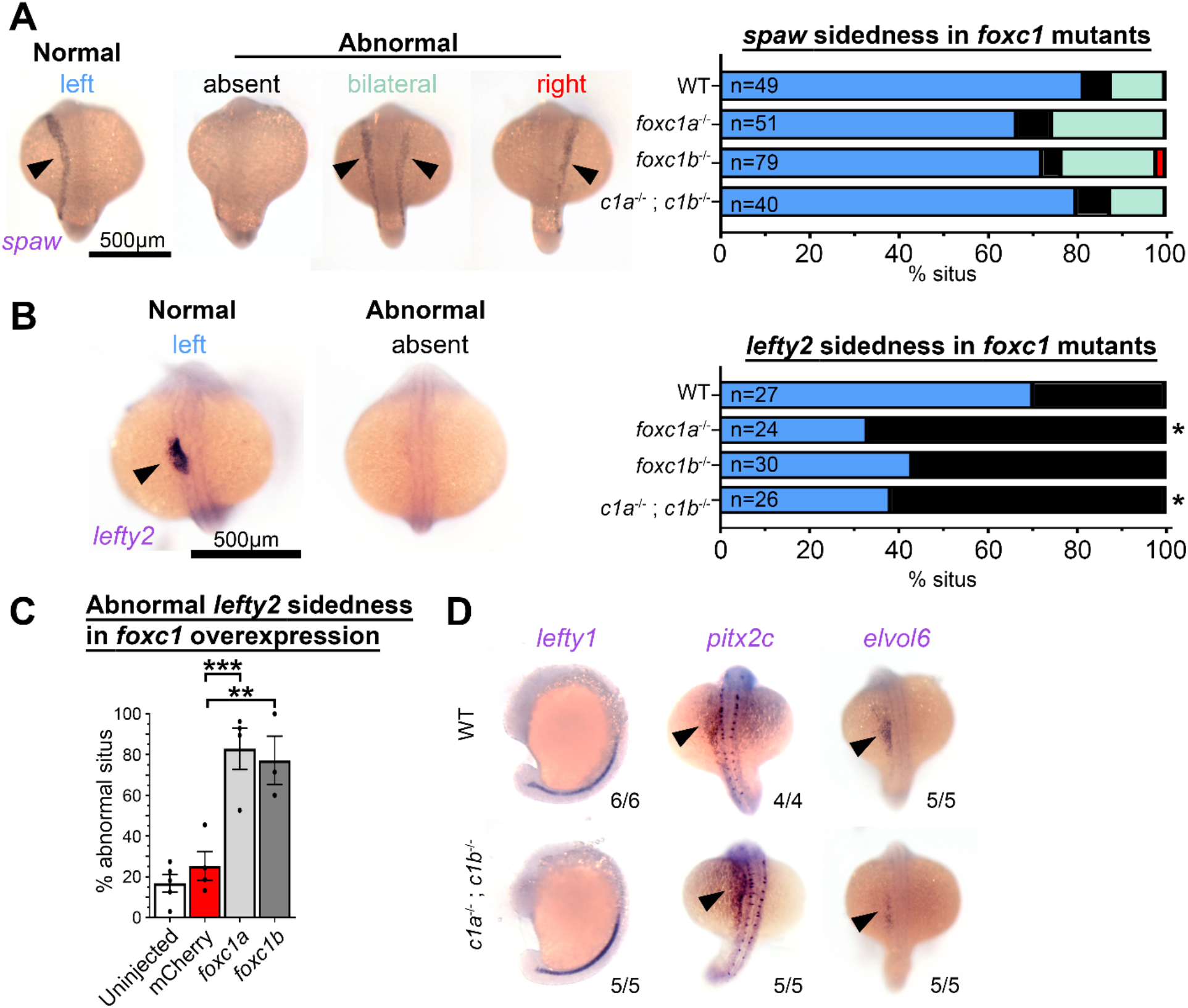
*foxc1* mutants display loss of *lefty2* expression independent of changes to other left-right patterning genes. (**A**) *In situ* hybridization with *southpaw* (*spaw*) revealed no difference in the prevalence of normal left-sided expression (blue) in *foxc1* mutants compared to controls (P>0.1, Fisher’s exact test). (**B**) Conversely *lefty2* expression was significantly altered in *foxc1* mutants. Normal left-sided *lefty2* expression (blue) was absent more frequently in *foxc1a*^-/-^ and double *foxc1a*^-/-^; *foxc1b*^-/-^ homozygotes (P= 0.012, 0.028, respectively), but unchanged in *foxc1b*^-/-^ homozygotes (p= 0.061 Fisher’s exact test). (**C**) *lefty2* expression was significantly abnormal with overexpression of *foxc1a* or *foxc1b* (P=0.0007 and 0.0030 respectively ANOVA and Dunnett Test). (**D**) Analysis of additional left-right patterning genes *lefty1, pitx2c*, and *elvol6* revealed no differences between WT and *foxc1a*^-/-^; *foxc1b*^-/-^ double homozygotes.

This finding of *lefty2* mis-expression led us to examine other genes required for establishment of the left-right axis. *lefty1* is a Spaw antagonist closely related to *lefty2*, but is expressed in the embryonic midline. *pitx2c* is asymmetrically expressed in the left LPM and lies downstream of *spaw*. No changes to expression of either gene were resolved in double homozygotes at 20 hpf (Fig. 6D). Finally, in situ analysis of *elovl6*, an enzyme asymmetrically expressed in lateral plate mesoderm that is responsive to LRO function (54, 55) revealed no changes in *foxc1a*^-/-^; *foxc1b*^-/-^ double homozygotes at 18 hpf. These results suggest that *foxc1* regulates *lefty2* rather than inducing loss or randomisation of all left-right patterning gene expression, as seen when LRO function is perturbed (56). Loss of *foxc1* may therefore be sufficient to perturb organ situs in a partially penetrant manner.

## Discussion

In this manuscript we demonstrate that CRISPR-Cas9 mutation of the zebrafish *foxc1a* and *foxc1b* paralogs recapitulates phenotypes observed in Axenfeld-Rieger syndrome (ARS) patients, and mouse models thereof (40, 41, 57). By mutating both zebrafish orthologs of human *FOXC1* we show that *foxc1a/b* homozygotes display craniofacial dysmorphism, cardiac defects and intracranial haemorrhage. These phenotypes correlate closely with clinical data, where up to 43% of patients exhibit craniofacial anomalies (23), 11% congenital heart defects (48) and 73% white matter hyperintensities (40). The absence of most phenotypes in *foxc1b*^-/-^ homozygotes suggests considerable genetic compensation. This concept is supported by *foxc1a*^-/-^; *foxc1b*^-/-^ double homozygotes but not *foxc1a* single mutants developing a high prevalence of hydrocephalus, which is a key phenotype of the murine *Foxc1*^-/-^ mutant *congenital hydrocephalus*, and an occasional finding in patients with heterozygous *FOXC1* mutation or deletion (58). From these data, we conclude that *foxc1a/b* mutant fish mirror several aspects of human and mouse ARS models.

The Forkhead box transcription factor family is evolutionarily conserved from yeast to humans and comprises more than 45 members in mammals, each containing an 80-100 amino acid winged-helix DNA-binding domain. Extensive links to development and disease have been established for many members of this family, and intriguingly, three Fox genes are recognized regulators of L-R patterning. *Foxj1* is expressed in the highly ciliated choroid plexus, lung epithelium, oviduct, and testis (59) and *Foxj1* knockout mice display loss of motile cilia and abrogation of left-right patterning. *Foxj1* is thus recognized as a key regulator of motile ciliogenesis (60). *Foxa2*, plays a role in promoting expression of *Pkd1l1*, and is essential for forming the left-right organizer (61), while *Foxh1* (also known as *Fast*) functions as a Smad co-factor downstream of Nodal signalling (62). Two additional findings led us to examine a role for Foxc1 in visceral organ situs. First, congenital heart defects (frequently associated with heterotaxy) are present in heterozygous patients and *Foxc1* knockout mice. Second, Pitx2, a protein that may directly bind Foxc1 (63) and and also causes ARS, is a critical regulator of left-right patterning. Our data demonstrate that *foxc1* mutation induces situs defects of visceral organs, with two mutant copies required for cardiac situs defects, while four are needed for anomalous gut situs. Recapitulation of these phenotypes with independently generated mutants and mRNA overexpression of either *foxc1a* or *foxc1b* demonstrates that increases and decreases in the levels of these *foxc1* paralogs impacts the situs of multiple visceral organs.

The establishment of left-right patterning can be divided into three separable phases: the creation of a ciliated epithelium that drives leftward fluid flow; the initiation of left-specific Nodal-Pitx2 gene expression; and finally the induction of a midline barrier that blocks Nodal activity from signaling on the right side. Our studies of Foxc1 demonstrate normal cilia-mediated flow in the LRO and proper initiation of left–specific *spaw* and *pitx2c*, consistent with Foxc1 playing a later role in regulating left-right axis formation. Indeed, when characterizing left-right isomerism defects in *foxc1a/b* mutants we note that phenotypes are not consistent with randomization of organ situs, as would have been expected from an early role in establishing left-specific gene expression. Furthermore, we find markedly perturbed *lefty2* expression in *foxc1a/b* mutants. Lefty proteins, which are known antagonists of Nodal (Spaw) signaling, have significantly greater rates of diffusion than the Nodal proteins that they antagonize (28). This leads to the prevailing model for how Lefty functions as a midline barrier to block extracellular Nodal from reaching the right side of the embryo. Consistent with the observed heterotaxic effects on the left-right placement of heart, liver, and pancreas, we conclude that Foxc1 proteins play a key role in the regulation of *lefty* gene expression, and are thus likely components of the establishment of the midline barrier for left-right patterning.

In this manuscript we present the first evidence of *foxc1* playing a role in left-right patterning of the lateral plate mesoderm, and in controlling organ situs. These data have significant implications for understanding the etiology of Axenfeld-Rieger syndrome associated congenital heart defects, and strengthen the case for cardiac screening in patients diagnosed with ARS. Our findings should also encourage a re-examination of organ situs in other Foxc1 models, since situs defects may be subtle and only partially penetrant. Since Foxc1 is the fourth Forkhead gene to participate in left-right patterning, this result emphasises the gene family’s importance in the control of organ situs, and will encourage future studies to determine how multiple Forkhead family members evolved seemingly distinct roles in the establishment of the left-right body axis.

## Materials and Methods

### Zebrafish lines and husbandry

Zebrafish lines were kept in accordance with the University of Alberta’s Animal Care and Use Committee guidelines. Animal care protocols were approved by the University of Alberta Biosciences Animal Care Committee with protocol number 00000082. Experiments were performed in the AB line, and *foxc1a*^UA1017^ and *foxc1b*^UA1018^ were generated and maintained on the AB background. The Tg(sox10:GFP)^ba4^ line was used for examination of the craniofacial deformities caused by *foxc1* mutation (64). Zebrafish embryos were raised at 28.5°C or 33°C in E3 media or E3 / 0.2mM 1-phenyl-2thiourea from 22hpf onwards to prevent pigmentation (65). Standard staging of embryos was conducted as in Kimmel et al. 1995 (66). *foxc1a^el542^*; *foxc1b^el620^* have been previously described (52).

### Bioinformatics

ENSEMBL sequences (Human ENSG00000054598, ENSG00000176692, mouse ENSMUST00000062292, ENSMUST00000054691, xenopus ENSXETG00000000594, ENSXETG00000016387, and zebrafish ENSDARG00000091481, ENSDARG00000055398) were aligned with Clustal Omega for sequence conservation analysis. The synteny of human and zebrafish *foxc1* orthologs were investigated in ENSEMBL using the “region in detail” options to explore the chromosomal regions surrounding each gene, and schematics were generated in Corel Draw.

### LRO cilia length measurements

Embryos raised at 33°c were collected at the 14 hpf and fixed overnight at 4°C in 4% PFA. Storage at −20°C in 100% methanol was usually performed for up to two weeks. Once transferred into PBS-T, embryos were dechorionated and blocked in 10% heat-inactivated goat serum, 1% bovine serum albumin for 1 hour while rocking. Primary antibody (monoclonal anti-tubulin, acetylated - Sigma T6793) at 1:1000 dilution was performed overnight at 4°C. Embryos were washed (3x 30 min PBS-T) and then secondary antibody (goat anti-mouse Alexa Fluor^®^ 555 - abcam ab150114) at 1:2000 and a nuclear counterstain (TO-PRO™-3 Iodide ThermoFisher Scientific T3605) at 1:2000 was performed for 2 hours before 3x 15 minute PBS-T washes and a final PBS-T wash overnight at 4°C while rocking. Embryos were transferred into 70% glycerol via gradient (30, 50, 70%) and the posterior region of the embryo including the LRO were excised with forceps and mounted in a slide viewing chamber and covered with a coverslip. The remainder of the embryo was processed for gDNA extraction. Imaging stacks through the whole LRO were performed on an LSM 700 confocal microscope (Zeiss) using a 40x oil lens with a 1μm Z-interval. Maximum intensity projections were generated and axonemal length for each cilium was calculated in FIJI, with the average cilia length per LRO per embryo being used for statistical analysis.

### Bead tracking

Left-right organiser function was assayed as described in (67). Embryos at the 11-12 hpf were dechorionated and embedded in a few drops of 1% low melting point agarose before the LRO was injected with approximately 0.5nL of Fluoresbrite Polychromatic Red 0.5 Micron Microspheres (Polysciences #19507). Bead flow was recorded for 10 seconds at 20x DIC on a Axioskop 2 microscope (Zeiss) using QCapture Plus. FIJI software using the Manual Tracker plug-in was used to produce the bead projections and record metrics.

### in situ hybridisation

*in situ* hybridisation was performed essentially as described in (68). Briefly, digoxigenin-labelled probes were synthesised from linear DNA templates using RNA polymerase and digoxigenin-UTP kits (Roche), and purified using SigmaSpin columns (Sigma). Fixed embryos were permeabilised with proteinase K treatment for 3 mins (18-20 hpf) or 15 mins (48 hpf), re-fixed with 4% PFA for 20 minutes, and hybridized with 1:200 RNA probes overnight at 65°c. High stringency washes in 0.2x and 0.1x SSC / 0.1% Tween 20 were carried out at 65°C for 20 minutes each, and blocking was performed for 1 hour in 2% sheep serum, 2 mg/ml bovine serum albumin. Anti-digoxigenin-AP antibody (Roche) at 1:5000 dilution was used to detect probe hybridisation, and NBT/BCIP (Roche) colouration reactions were performed at 33°C until signal was saturated. In situs were imaged by dissection light microscopy and then the tissue was genotyped. The probes used were *myl7* (69), *foxa3* (70), *pitx2c* and *elvol6* (54), *foxc1a* and *foxc1b* (51), *spaw* fwd: ATGCAGCCGGTCATAGC, rev: TCAATGACAGCCGCACTC, *lefty1* fwd: ATATTCTGACACGACACGTC, rev: CTGAAATATTGTCCATTGC, *lefty2* fwd: ATCAAGTACTCGGACACC, rev: GGAGTCCCATAACTGTG.

### Light Microscopy

Live imaging was performed on PTU-treated embryos anaesthetised in 0.6mM Tricaine (Sigma Cat.# A-5040) as per (71). Both live imaging and *in situ* processed embryos were imaged on a 1% agarose coated dishes using a SZX12 light microscope (Olympus) with QCapture Suite Plus. White balance and brightness / contrast editing was performed in Adobe^®^ Photoshop^®^, and was performed consistently between all genotypes and conditions.

### mRNA overexpression and morpholino microinjection

Capped mRNA was generated from a linear DNA template in a pCS2+ vector using the mMessage mMachine (ThermoFisher Scientific) kit, purified using TRIzol™ (ThermoFisher Scientific) and microinjected into the one-cell stage embryo at doses of 5, 15 and 75 pg. The highest 75 pg dose was used for all subsequent experiments unless otherwise stated. All RNA technical replicates were performed on a single day, injecting *foxc1* transcripts and control RNA, at the same dose, into the same clutch of embryos. Morpholino oligonucleotides *foxc1a* – CCTGCATGACTGCTCTCCAAAACGG, *foxc1b* – GCATCGTACCCCTTTCTTCGGTACA were previously reported (40).

### Organ situs scoring

Cardiac situs scoring was performed by light microscopy on live 48hpf embryos. While still in the chorion, embryos were anaesthetised and rolled with fine watchmaker’s forceps so they could be viewed ventrally. The sequential atrial-ventricular heart contractions allowed for the cardiac situs to be easily scored based on looping of the ventricle in comparison to the atrium. To assess gut situs, *in situ* hybridisation on 48hpf embryos was performed using a *foxa3* probe and once developed, images of each embryo were recorded. Scoring of heart and gut situs was performed in a masked manner and subsequently tissue was processed for gDNA extraction.

### Genotyping

Embryonic tissue was dissociated in 50mM NaOH as in (72) and then diluted 1:10 before being used as template for PCR. Both *foxc1* mutations were resolved via standard PCR genotyping: *foxc1a fwd*: TTCTTCGCCAGCTGTACG, rev: AATAACTTTGGTCGCTGC, *foxc1b* fwd: CCGTGTCTAGCCAAAGC, rev: TCGGATGAGTTTTGGATG. Gel electrophoresis was used to resolve the WT and mutant bands under the following conditions: *foxc1a* – SB buffer, 3% agarose, 300V for 1 hour (as in (73)), *foxc1b* – TAE buffer, 2% agarose, 150V for 1 hour.

### Statistical analysis

Situs scoring of the heart and gut from *foxc1* mutants were categorical data so Fisher’s exact test was performed comparing normal and abnormal frequency between mutants and the WT control. Embryos from at least five clutches of embryos were scored and pooled for our analysis. For both mRNA overexpression and morpholino knockdown the percentage with situs abnormalities was calculated per technical replicate (a clutch of embryos injected with the mRNA or MO). The mean percentage situs was compared via ANOVA with Dunnett post-hoc testing to determine which categories were significantly different. All graphs and statistics were performed in GraphPad Prism 8.

## Acknowledgements

Funding was provided by the Canadian Institutes of Health Research, and Women and Children’s Health Research Institute (to OJL), a Heart and Stroke Foundation of Canada postdoctoral fellowship (to CRF), by the Natural Sciences and Engineering Research Council of Canada (NSERC, RGPIN 2016-06482 to AJW), NIH grant DE027550 to JGC and by Bridge funding from Women & Children’s Health Research Institute (to AJW).

The authors thank Prof. Jeffrey Amack for providing in situ hybridisation probe templates, and for his insight into the results early in the project; Dr. Sarah Hughes for access to her imaging suite; Casey Carlisle, Dr. Lance Doucette, and Nicole Noel for critical reading of the manuscript; and Science Animal Support Services technicians at the University of Alberta.

---

**Table S1.**
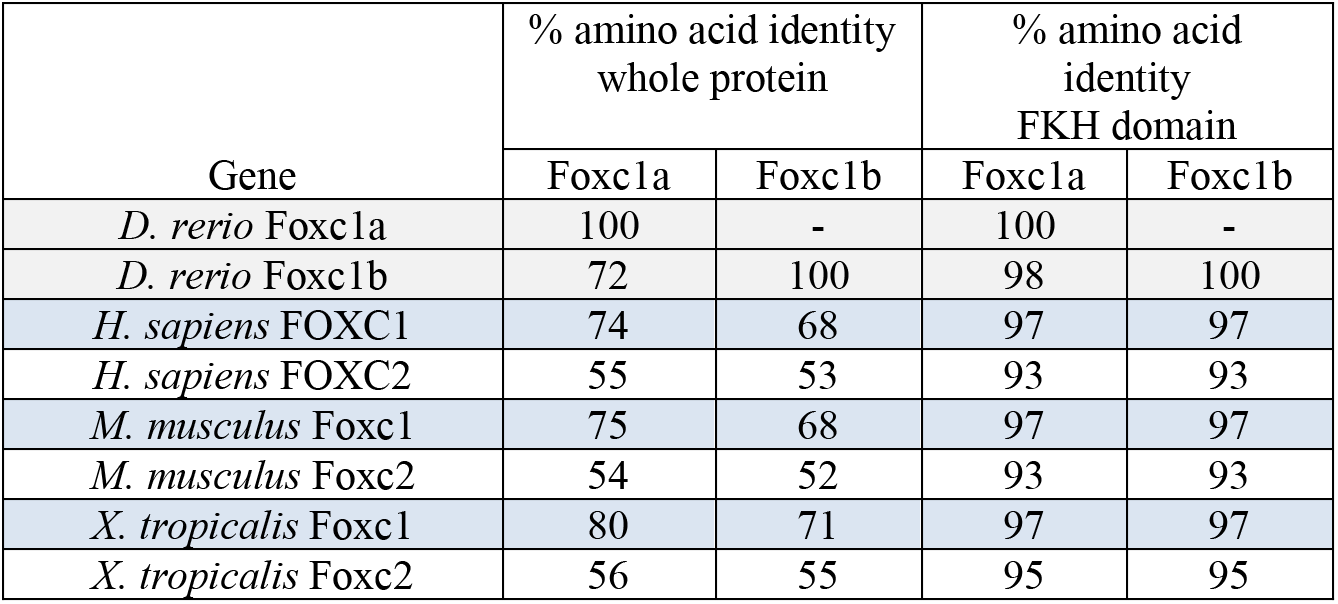
Both zebrafish Foxc1 protein sequences are orthologous to human FOXC1. The zebrafish Foxc1a and Foxc1b protein sequences were aligned against each other (grey) and the human, mouse and frog FOXC1 (blue) and FOXC2 (white) sequences. Irrespective of whether the whole protein or the *Forkhead* domain (FKH) were compared, the zebrafish paralogs were more similar in sequence to FOXC1 orthologs than FOXC2.

**Fig. S1.**
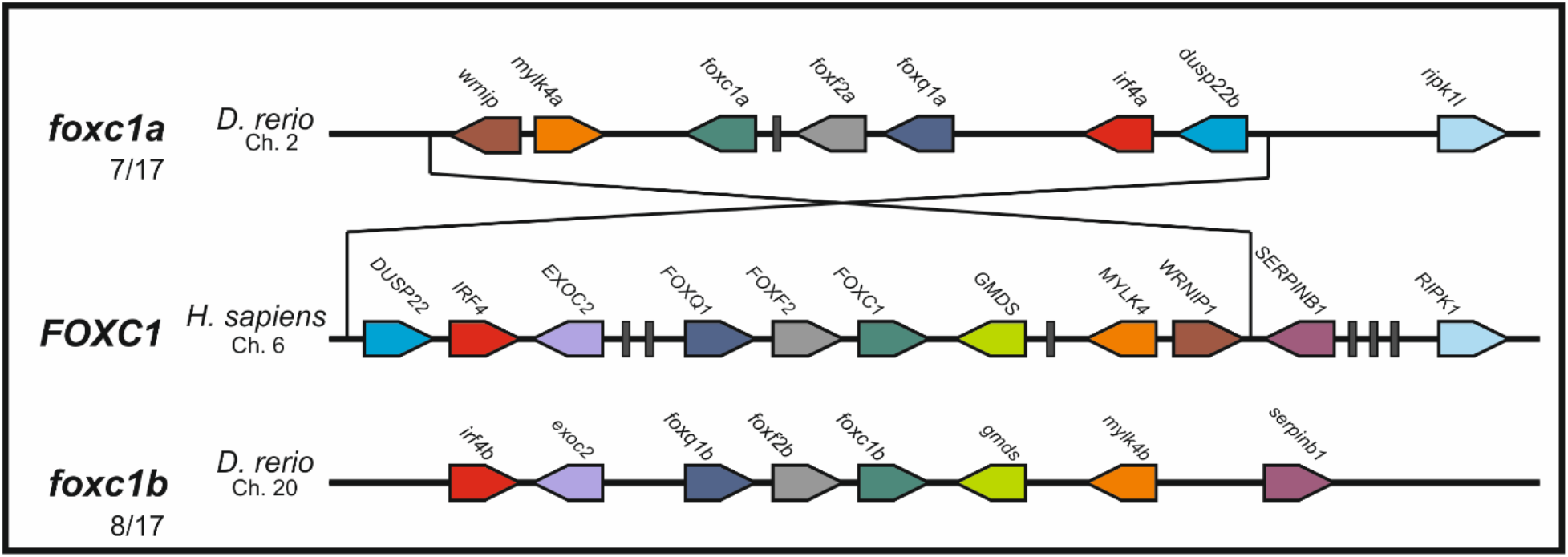
Syntenic analysis of the *foxc1* paralogs. Schematic representation of the chromosomal regions surrounding human *FOXC1*, and zebrafish paralogs. *foxc1a* shares 7 of 17, and *foxc1b* 8 of 17, orthologous genes with the corresponding human chromosomal region and both zebrafish chromosomes maintain the conserved *foxq1, foxf2, foxc1* triplet.

**Fig. S2.**
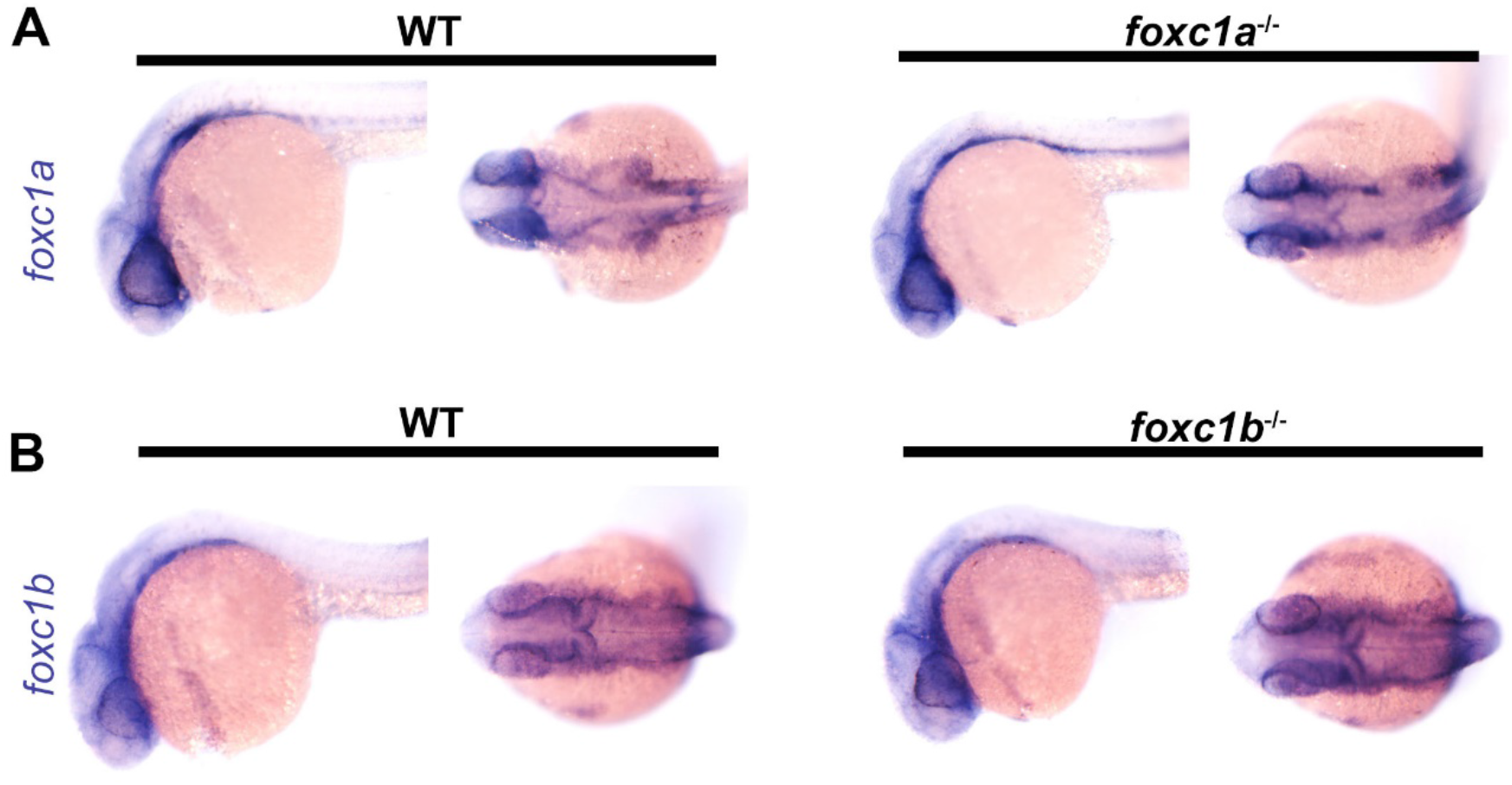
*foxc1* mutations *foxc1a*^ua1017^ and *foxc1b*^ua1018^ does not induce nonsense mediated decay. In situ hybridisation at 24 hpf using *foxc1a* (**A**) and *foxc1b* (**B**) specific probes demonstrates that mutant mRNA is not degraded in *foxc1a*^ua1017^ or *foxc1b*^ua1018^ homozygotes respectively.

**Fig. S2.**
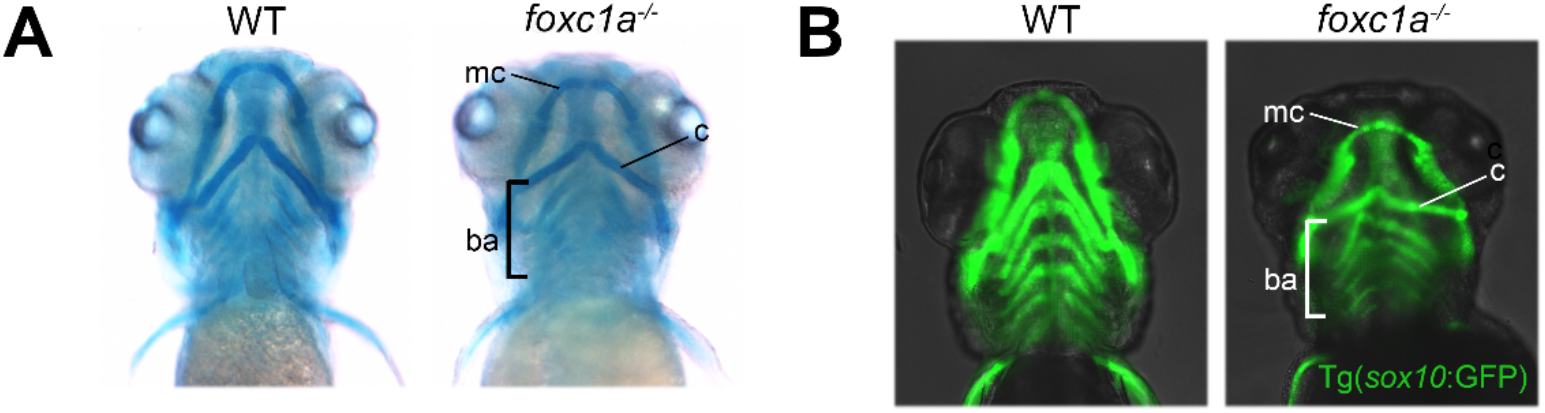
*foxc1a* mutants have defects in craniofacial cartilage. (A) Alcian blue staining of the craniofacial skeleton at 5 dpf. *foxc1a*^-/-^ homozygotes display craniofacial dysmorphism compared to WT controls. (**B**) The *Tg*(*sox10:GFP*) transgene remains visible at 5 dpf in the craniofacial skeleton and reveals disorganisation of the brachial arches (ba), as well as a significant reduction in the size of Meckel’s (mc) and the Ceratohyal cartilages (c)

**Fig. S4.**
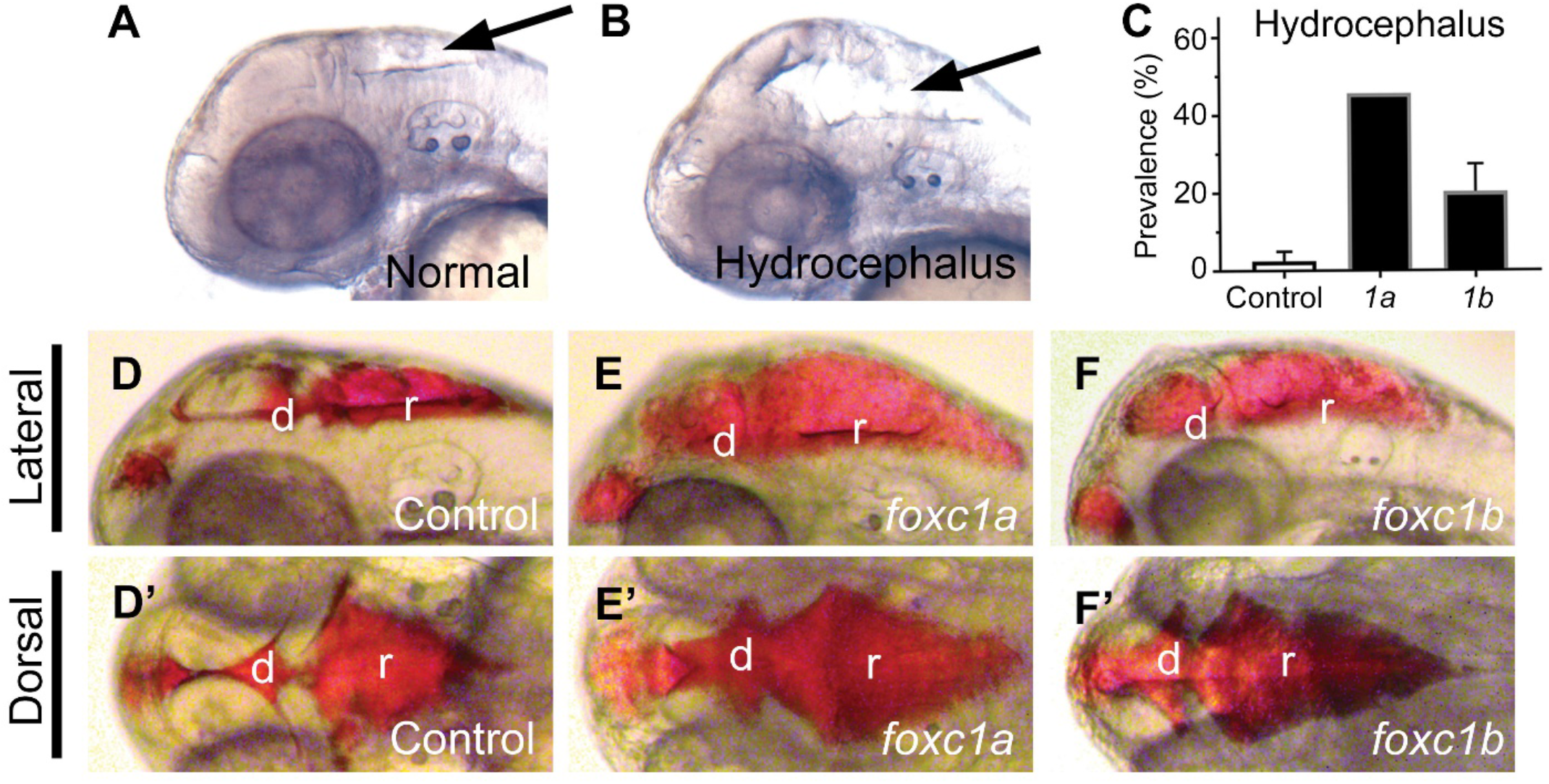
Overexpression of *foxc1a* or *foxc1b* causes hydrocephalus. Compared to 75 pg mCherry injected controls (**A**), 75 pg of *foxc1a* or *foxc1b* mRNA result in development of hydrocephalus (arrow) at 72 hpf (**B**) at a prevalence of 46% and 20%, respectively (**C**). Microinjection of rhodamine-conjugated dextran into the diencephalic ventricle allows for visualisation of the entire cerebral ventricle system (**D-F’**). In both *foxc1a* and *foxc1b* OE there is expansion of the diencephalic (d) and rhombencephalic (r) ventricles. Panels show hydrocephalic cerebral ventricles from both lateral (**D-F**) and dorsal (**D’-F’**) aspects.

**Fig. S5.**
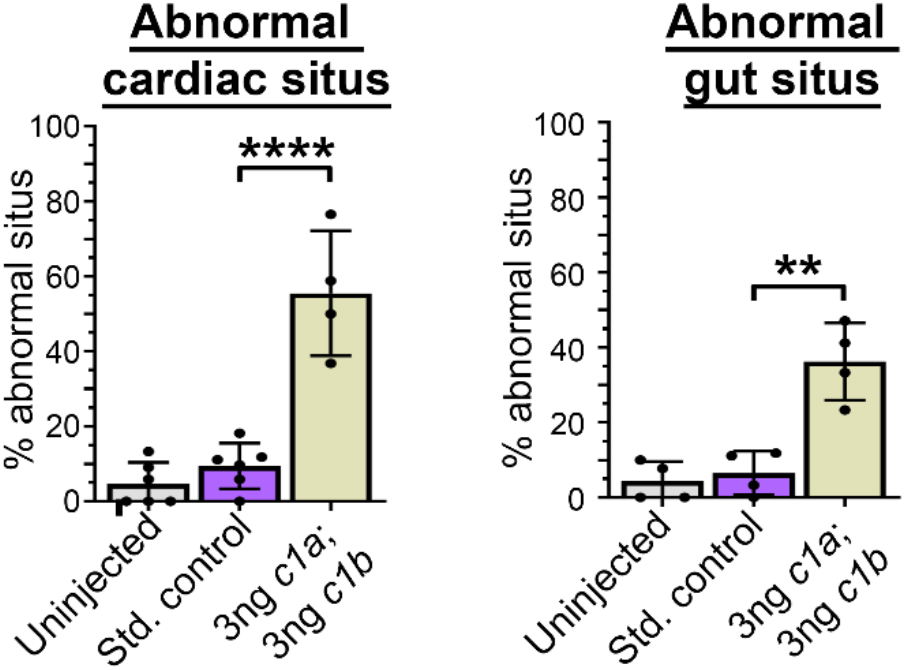
Morpholino knockdown recapitulates mutant situs phenotypes. Abnormal organ situs was significantly more frequent when *foxc1a* and *foxc1b* were knocked down via morpholino oligonucleotide (cardiac: P<0.0001; gut, P= 0.0003, ANOVA and Dunnett test).

**Fig. S6.**
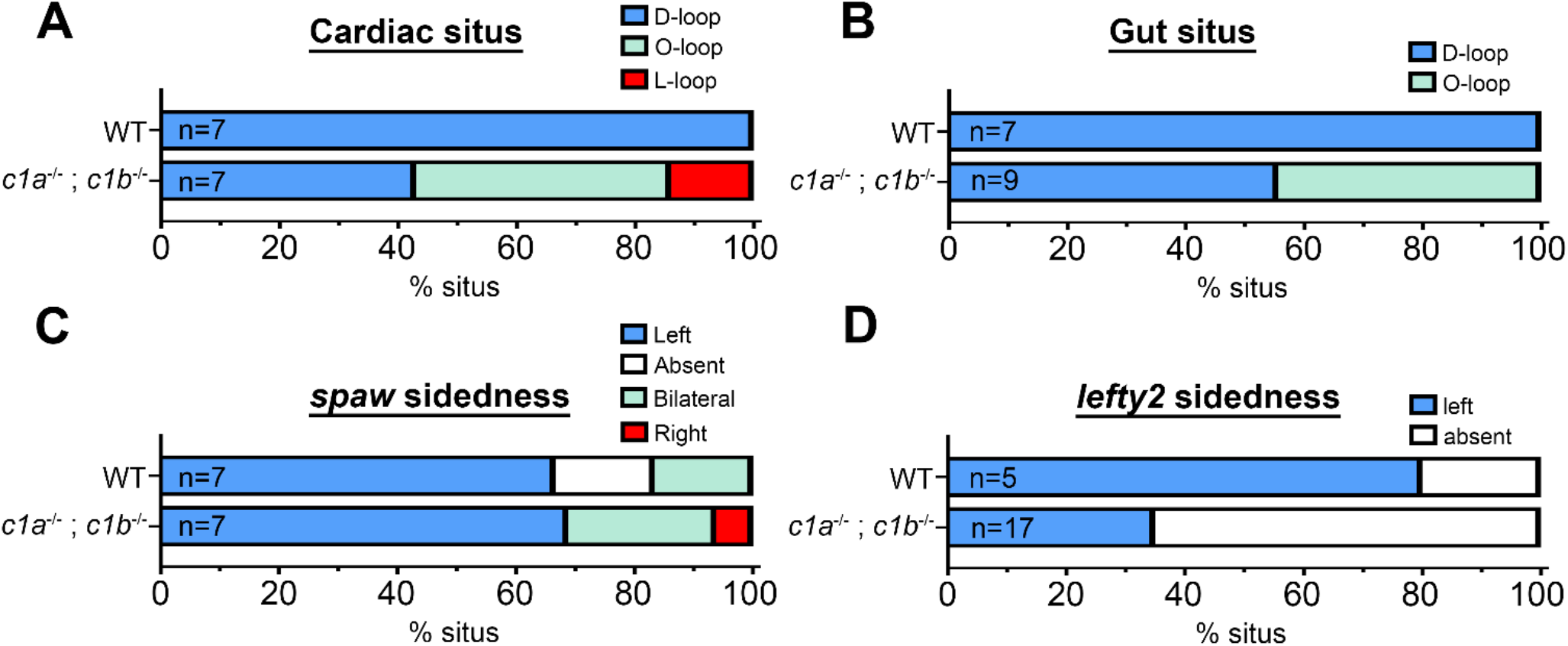
Organ situs and left-right patterning defects were observed in distinct alleles of *foxc1a*^-/-^; *foxc1b*^-/-^. Examination of *foxc1a^el542^; foxc1b^el620^* double homozygous mutants at 48 hpf revealed both cardiac situs (**A**) and gut situs (**B**) defects. At 18 hpf sidedness of *southpaw* (**C**) was comparable between WT and double mutants (P>0.99), however *lefty2* expression (**D**) was more frequently absent in double mutants compared to WT.

